# Directional coordination along the cortical hierarchy is selectively reorganized in schizophrenia

**DOI:** 10.64898/2026.05.25.727703

**Authors:** Sir-Lord Wiafe, Najme Soleimani, Zening Fu, Robyn Miller, Vince D. Calhoun

**Affiliations:** Tri-Institutional Center for Translational Research in Neuroimaging and Data Science (TReNDS), Georgia State University, Georgia Institute of Technology, and Emory University, Atlanta, GA 30303, USA

## Abstract

Schizophrenia has long been framed as a disorder of the cortical hierarchy. Yet, the hierarchy has been studied almost entirely as fixed anatomy, leaving its moment-to-moment expression in ongoing brain activity unmeasured. By retaining the signed lead-lag geometry that conventional connectivity measures discard, complex-valued phase synchrony resolved a dynamic directional axis of coordination in which regions reversed their lead-lag relationship over time as the brain transitioned among feedback, feedforward, and integrative regimes. The three regimes replicated across seven datasets from four independent cohorts and aligned region by region with an independently defined cortical hierarchy. Neighboring organizing principles, including propagating waves, arousal, and the principal gradient, were insufficient to explain regime identity. Within the same subjects, GABAergic sedation shifted coordination toward feedback and motor imagery toward feedforward, demonstrating that the axis reflects a manipulable property of brain dynamics rather than a statistical partition. In schizophrenia (151 patients, 160 controls), the axis reorganized selectively: feedback coordination became less persistent, integrative coordination more persistent, feedforward comparatively spared, and trajectories less diverse. These effects replicated in an independent clinical cohort (68 patients, 89 controls). The reorganization tracked positive symptom severity, and its relationship to cognition reversed relative to controls: faster convergence to equilibrium was associated with better cognition in controls and worse cognition in patients. Directional coordination therefore identifies a reproducible and controllable dynamic expression of the cortical hierarchy that is selectively reorganized in schizophrenia and linked to both symptoms and cognitive function.

## 1. INTRODUCTION

The most influential computational accounts of schizophrenia are theories of direction, centered on the balance of ascending and descending signaling in the cortical hierarchy. The dysconnection hypothesis^1,2^ and predictive-coding models^3,4^ locate the disorder in a failure of descending signals to constrain ascending ones, so that sensory evidence escapes the predictions that should contextualize it. In computational terms, the disorder reflects aberrant precision, the weighting that determines how far ascending signals update descending models. On these accounts, positive symptoms emerge when this balance fails^5–8^. That balance has been examined only indirectly, through local electrophysiological signatures of prediction^9,10^, or within small preselected circuits^11,12^, while the measures that dominate large-scale studies of schizophrenia^13,14^ carry no directional information.

The hierarchy itself is well established anatomically and functionally. Anatomical tracing defined the ascending and descending pathways of primate cortex^15,16^, electrophysiology assigned them distinct oscillatory channels^17^, and macroscale proxies place every region along a sensory-to-association axis, from the principal connectivity gradient^18^ to gradients of intrinsic timescale^19,20^. Spontaneous activity traverses this axis as propagating waves^21,22^. In schizophrenia, this architecture is compressed, with reduced separation between sensory and association systems^23,24^, and its dynamic markers shift with it. Hierarchical gradients of intrinsic timescales are altered in psychosis in a symptom-specific manner^25^, and electrophysiological signatures of hierarchical prediction, from corollary discharge to mismatch negativity, are attenuated in patients, representing some of the most replicated biological findings in the disorder^9,10^. In parallel, resting-state fMRI established that coordination is dynamic, moving through recurring connectivity states whose temporal properties are altered in the disorder^13,14,26,27^, and meta-state analyses showed that patients traverse a narrower and less dynamic range of connectivity configurations^28,29^. These studies establish where regions sit along the hierarchy, how strongly they couple over time, and that prediction is disturbed in patients. What they do not establish is which level of the hierarchy leads at a given moment.

The omission is built into the measures. Functional and dynamic functional connectivity quantify the strength of synchrony and typically discard directional phase information, collapsing opposite lead-lag configurations into similar coupling strength^30–33^, and phase-locking state analyses inherit the same convention^34,35^. For instance, two regions phase-locked at plus and minus sixty degrees are identical to these measures, although which region leads is reversed. Edge-centric and co-fluctuation approaches resolve coordination at finer temporal grain, yet they too are built on undirected products^36,37^. Lag analyses of resting fMRI recovered reproducible temporal orderings but averaged them over entire scans, potentially merging configurations of opposite direction^38^. Lag structure is itself altered in schizophrenia, though these analyses index timing offsets and do not resolve which level of the hierarchy leads^39^. Directed models such as Granger causality and dynamic causal modeling estimate influence explicitly but under strong model assumptions and are generally applied to small, preselected networks^11,40^, limiting their ability to discover whole-brain directional organization without specifying its form in advance. Decades after the hierarchy was mapped, a directional theory of schizophrenia is still being tested, at the whole-brain level, with directionless measurements.

Here we retain the quantity these measures discard. Using complex-valued phase synchrony^41^, which preserves the signed lead-lag geometry of interregional phase relationships, we sought a whole-brain measure of this directional balance. We asked whether ongoing coordination is directionally organized, whether that organization can be moved experimentally, and whether it is disrupted in schizophrenia. Across seven resting-state datasets from four independent cohorts, coordination moved through three recurring regimes: feedback with association cortex leading, feedforward with sensory cortex leading, and integrative with balanced exchange. The discovery cohort was fBIRN, and replication spanned COBRE, four sessions of HCP-Aging, and HCP-Adult. The regimes were identified from the phase geometry alone, without anatomical labels or clinical variables, and the ordering of regions within the directional regimes aligned with an independently defined cortical hierarchy^18^.

This directional measurement revealed a selective reorganization in schizophrenia. The hierarchical failure was not symmetric across directions. Feedback coordination was less persistent while feedforward coordination was comparatively spared, with dynamics shifting towards the integrative regime that carries little net directionality. These findings replicated in an independent clinical cohort. The reorganization also carried opposite cognitive associations across groups, faster convergence to equilibrium was associated with better verbal learning in controls but with worse attention and composite cognition in patients. This sign reversal is difficult to reconcile with a simple global weakening of connectivity, and the patient associations remained unchanged with antipsychotic dose as a covariate. Positive symptoms were associated with a shift away from balanced exchange and toward feedforward coordination. Critically, the directional axis was experimentally manipulable. Within the same healthy subjects, sedation with propofol, a GABA-A positive allosteric modulator^42^, shifts coordination toward feedback while motor imagery shifts it toward feedforward, placing the disorder’s abnormality on a dimension under experimental control. Control analyses show that the directional structure is not attributable to a fixed vascular delay and is not recoverable from propagating waves, arousal, or the principal gradient^21,43^. Directional coordination thus provides a whole-brain dynamic measure directly aligned with hierarchical accounts of schizophrenia and reveals a selective disruption of directional organization that conventional undirected measures cannot capture.

## 2. MATERIALS AND METHODS

### 2.1. Complex-valued phase synchrony for directional coordination

We quantified directional coordination using complex-valued phase synchrony (CVPS)^41^, which preserves both the strength and the sign of interregional phase relationships. Let *x*(*t*) be the BOLD time series at one region, sampled at the repetition time (TR). Its instantaneous phase *φ_x_*(*t*) was estimated by convolving *x*(*t*) with a complex Gabor wavelet that is both frequency- and time-localized:

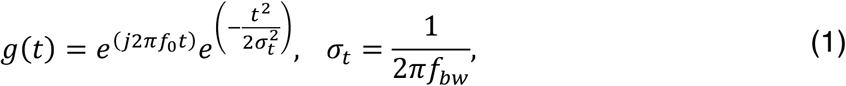

where *f*_0_ = 0.05 *Hz* is the center frequency, and *f_b_*_w_ = 0.02 *Hz* is the half-bandwidth, matching the frequency range conventional in fMRI phase-synchrony studies^31–33^. Wavelet estimation was chosen over the more common Hilbert approach because it tracks phase more accurately under non-stationary BOLD-like conditions (Fig. SF. 1e). For two regions with instantaneous phases *φ_x_*(*t*) and *φ_y_*(*t*), the asymmetric complex phase synchrony signal was defined as

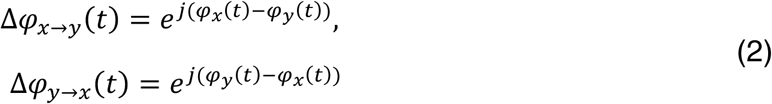

whose magnitude reflects synchrony strength and whose angle carries the signed lead-lag between the regions. We refer to this signed lead-lag dimension as directional coordination. Kernel truncation and normalization, the analytic-signal derivation, and the relation of CVPS to the dynamic co-modulation framework^44^ are provided in the Supplementary Methods (“Complex-valued phase synchrony mathematics,” Eqns. SE. 1 and SE. 2). Simulations demonstrating that cosine-based measures cannot separate opposite lead-lag configurations, and empirical illustrations of the information carried by the preserved sign, are provided in the Supplementary Methods (“Simulation studies”) and Fig. SF. 1.

### 2.2. fMRI data & processing

We analyzed resting-state fMRI from Phase III of the Function Biomedical Informatics Research Network (fBIRN) consortium^45^ as the discovery cohort, comprising 160 healthy controls (37.0 ± 10.9 years, 45 F/115 M) and 151 individuals with schizophrenia (38.8 ± 11.6 years, 36 F/115 M), scanned at seven sites with a repetition time of 2 s. Data underwent standard preprocessing (discarding of initial volumes, slice-timing correction, rigid-body realignment, MNI normalization, 6 mm smoothing)^46^, and 53 intrinsic connectivity networks (ICNs) were extracted per subject using the fully automated, spatially constrained NeuroMark ICA pipeline with the neuromark_fMRI_1.0 template^47^. ICN time courses were z-scored before CVPS analysis. Replication used three independent cohorts processed with the identical NeuroMark framework, COBRE^48^ (n = 157, comprising 68 individuals with schizophrenia and 89 controls, single site), four sessions of HCP-Aging^49^ (n = 718 per session), and HCP-Adult^50^ (n = 827). Full demographic and acquisition details for all cohorts are provided in the Supplementary Methods (“Supplementary fMRI data”). All data were collected in the source studies under institutional review board approval with informed consent and were analyzed retrospectively here.

### 2.3. Task and anesthesia cohort

To test whether the directional regimes generalize beyond the resting state and the ICA-derived parcellation, and whether the axis can be moved experimentally, we analyzed 26 healthy volunteers (OpenNeuro ds006623) scanned under two within-subject manipulations^51^, propofol sedation titrated to loss of responsiveness (unconscious vs conscious) and motor and spatial imagery (the Owen paradigm, imagining playing tennis or navigating one’s home; imagery vs rest)^52^. We analyzed the preprocessed time series distributed with the dataset (fMRIPrep with XCP-D denoising), parcellated with the Glasser-360 atlas^53^. Regions were grouped into sensory and association systems using the Yeo network labels provided with the dataset for computation of the DHI. Regimes were re-derived by clustering within this cohort, so recovery of the same three regimes constitutes independent replication and not imposed structure. Within-subject differences in regime occupancy were tested with two-tailed paired t-tests, FDR-corrected, with effect size as paired Cohen’s dz computed as manipulation minus baseline, so that positive values indicate increased occupancy under the manipulation. Acquisition parameters, propofol titration details, and task timing are provided in the Supplementary Methods (“Task and anesthesia cohort”).

### 2.4. Identifying recurring directional coordination regimes

Coordination dynamics were modeled as a sequence of discrete regimes, consistent with prior work representing time-resolved fMRI dynamics as transitions among a finite set of states^27^. Each complex edge value was represented as a two-dimensional real vector of its cosine and sine components, retaining one direction per pair, an embedding that preserves the circular geometry of phase relationships (Supplementary Methods, “K-means distance selection,” Eqns. SE. 3 and SE. 4). Regimes were identified by k-means clustering with the city-block distance (20 initializations), and the number of clusters was set to three by the elbow criterion. The regime architecture did not depend on this choice, as the integrative regime was present at every K and the two directional poles separated cleanly at K = 3, with K = 4 only subdividing the integrative regime (Supplementary Fig. SF. 2). Cluster centroids were mapped back to the complex domain to yield representative coordination patterns per regime.

### 2.5. Vascular and non-neural control analyses

A directed lead-lag could in principle reflect a fixed, non-neural hemodynamic transit delay. Two control analyses, run identically across all seven datasets, tested this account using the cosine (CRP) and sine (SRP) of the relative phase between each pair of the 53 ICNs, which index undirected coupling and signed lead-lag, respectively. First, we compared the CRP and SRP patterns between the two most directionally opposite regimes across the 1,378 pairs, at the group and single-subject level, and computed the percentage of pairs whose lead-lag sign reversed within a scan, quantities that test whether direction is fixed across regimes. Second, we recomputed the group-mean signed relative phase in six narrow sub-bands spanning 0.02 to 0.13 Hz and derived directional strength, the implied delay, and the directional pattern per cohort and frequency, quantities that test whether the phase offset behaves as a fixed conduction delay. Sub-band definitions, per-cohort preprocessing, and scan-length considerations are provided in the Supplementary Methods (“Vascular and non-neural control details”).

### 2.6. Quantifying hierarchical directional flow within coordination regimes

To quantify the dominant direction of coordination flow within each regime, each cluster centroid was represented as a complex-valued edge vector, with the value of edge *e* in regime *k* written as *Z* = *Re*^jΔ*φ*^, where Δ*φ* carries the direction of the lead-lag relationship and the magnitude *R* carries its reliability. To identify the edges that most strongly distinguish a regime from the others, each edge received a discriminative score:

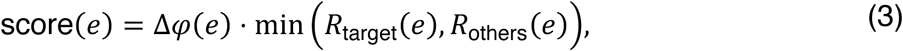

where Δ*φ*(*e*) denotes the minimum circular difference between two phase angles and the minimum is taken over the edge magnitudes across regimes, so that the score is high only for edges showing both strong directional separation between regimes and reliable phase structure. Node-level driver scores were then computed by aggregating signed edge contributions across these scores, with the direction of each edge’s influence given by the sign of the phase difference between its two nodes (Supplementary Methods, “Directional driver scores”). Nodes were grouped into sensory (visual, sensorimotor, auditory) and association (cognitive control, default mode) systems; subcortical and cerebellar networks were excluded from the index. The directional hierarchy index (DHI) was defined as the difference in mean driver score between association and sensory systems, so that positive values indicate association-leading flow and negative values indicate sensory-leading flow. Departure from zero was tested against a null distribution generated from 50,000 random system-label permutations.

### 2.7. Testing alignment with an externally defined cortical hierarchy

To test whether the directional ordering recovered by CVPS reflects an independently defined cortical hierarchy, we correlated each region’s net directional driver within a regime against its position on the principal functional-connectivity gradient^18^, computed by diffusion-map embedding (BrainSpace^54^) from the group functional connectivity of 207 unrelated healthy HCP adults distributed with the ENIGMA toolbox^55^. Because this gradient derives from undirected connectivity in separate subjects, and neither our lead-lag measures nor our sensory-association grouping enters its construction, any correspondence is non-circular with respect to how the regimes are defined. Driver scores were delta-weighted to down-weight edges whose relative phase is shared across regimes, so that the correlation reflects regime-specific reorganization. Spearman correlations spanned 42 cortical components in fBIRN and 329 cortical regions in the manipulation cohort, with significance assessed against spatial-autocorrelation-preserving nulls generated by Moran spectral randomization (1,000 surrogates, two-sided)^54,56^. Projection of the gradient into the NeuroMark component space, robustness to the gradient kernel, and a convergent cytoarchitectonic check are described in the Supplementary Methods (“External hierarchy alignment”).

### 2.8. Testing whether the directional regime is recoverable from neighboring principles

To assess whether the feedforward-versus-feedback distinction is contained in neighboring organizing principles, we tested how well each construct could decode the active directional regime from moment to moment, independently in all seven datasets. Each time point was assigned to a regime by nearest centroid; the two directional regimes were retained as a binary label, and up to 30 time points per regime per subject were sampled so that no subject or over-occupied regime could dominate. From the same time points, we extracted five feature sets: the CVPS lead-lag features themselves, which serve as a reference upper bound since the regimes are defined in that space, and four competing constructs, propagating-wave patterns^21^, propagating-wave phase^22^, the global/arousal signal^43^, and the principal gradient projection^18^. For each feature set, an L2-regularized logistic regression with balanced class weights was trained under subject-grouped five-fold cross-validation, so that no subject appeared in both training and test folds, and performance was summarized as balanced accuracy (chance = 0.5), with uncertainty from 2,000 bootstrap resamples over subjects. Identical pipelines, folds, and sampling were applied to every feature set, so that the feature set was the only quantity that varied. Feature-set construction details are provided in the Supplementary Methods (“Decoding feature sets”).

### 2.9. Characterizing coordination dynamics using Markov chain properties

The sequence of regime assignments was modeled per subject as a first-order Markov process over the three regimes. Regime persistence was quantified as mean dwell time, the average duration spent in a regime once entered, and occupancy as fraction rate, the proportion of time spent in each regime^57^. Subject-specific transition probability matrices yielded the stationary distribution, the long-run equilibrium over regimes^57^; the spectral gap, one minus the second-largest eigenvalue, indexing the rate of convergence to equilibrium; the mixing time, the number of steps required to approach the stationary distribution within a fixed tolerance; and the entropy rate, the uncertainty of regime transitions, reported as a percentage of the maximum for a three-state system. Formal definitions and derivations are provided in Wiafe et al.^58^ and the Supplementary Methods (“Markov chain metrics, formal definitions”).

### 2.10. Constructing a persistence landscape of directional coordination dynamics

To visualize the global organization of coordination dynamics, each regime’s stability was defined as the product of its stationary probability and its self-transition probability, capturing how often it is visited and how persistently it is maintained. Regimes were embedded on a two-dimensional simplex, and a continuous landscape was generated by summing isotropic Gaussian kernels with amplitudes proportional to stability, inverted so that deeper basins correspond to more stable regimes. Group landscapes were scaled to a common reference so that basin-depth differences reflect relative regime stability. Kernel and scaling details are provided in the Supplementary Methods (“Persistence landscape construction”).

### 2.11. Statistical framework for clinical and cognitive association

Group differences between healthy controls and individuals with schizophrenia were assessed using generalized linear models with age, sex, scanning site, and mean framewise displacement as covariates, applied to all coordination metrics (mean dwell time, fraction rate, transition probabilities, stationary distribution, spectral gap, mixing time, and entropy rate). Identical models were applied in COBRE, omitting the site covariate because COBRE is a single-site cohort. Multiple comparisons were controlled using the Benjamini-Hochberg false discovery rate, applied across the full family of coordination metrics within each cohort, and an effect was considered replicated when it was significant after correction in both cohorts with the same direction. Within-group associations with cognitive performance, measured with the Computerized Multiphasic Interactive Neurocognitive System (CMINDS), and with Positive and Negative Syndrome Scale (PANSS) scores were assessed with the same models and covariates, conducted separately within each group and FDR-corrected within each family of tests (transitions, fraction rate metrics, and global dynamical metrics corrected as separate families). As a sensitivity analysis, all within-patient association models were re-estimated with chlorpromazine-equivalent dose as an additional covariate; results were unchanged (Supplementary Table ST. 1).

## 3. RESULTS

### 3.1. Ongoing brain activity resolves into three directional coordination regimes that recur across independent cohorts

Applying complex-valued phase synchrony to whole-brain resting-state activity in the discovery cohort (fBIRN, n = 311), we found that directional coordination evolved along a structured axis comprising three recurring regimes: a feedback-dominated regime, a feedforward-dominated regime, and an integrative regime of balanced bidirectional exchange. To quantify the dominant direction of flow within each regime, we computed a directional hierarchy index (DHI), the net lead-lag between association and sensory systems, with positive values indicating association-leading flow and negative values indicating sensory-leading flow (Fig. 1a).

**Fig. 1:**
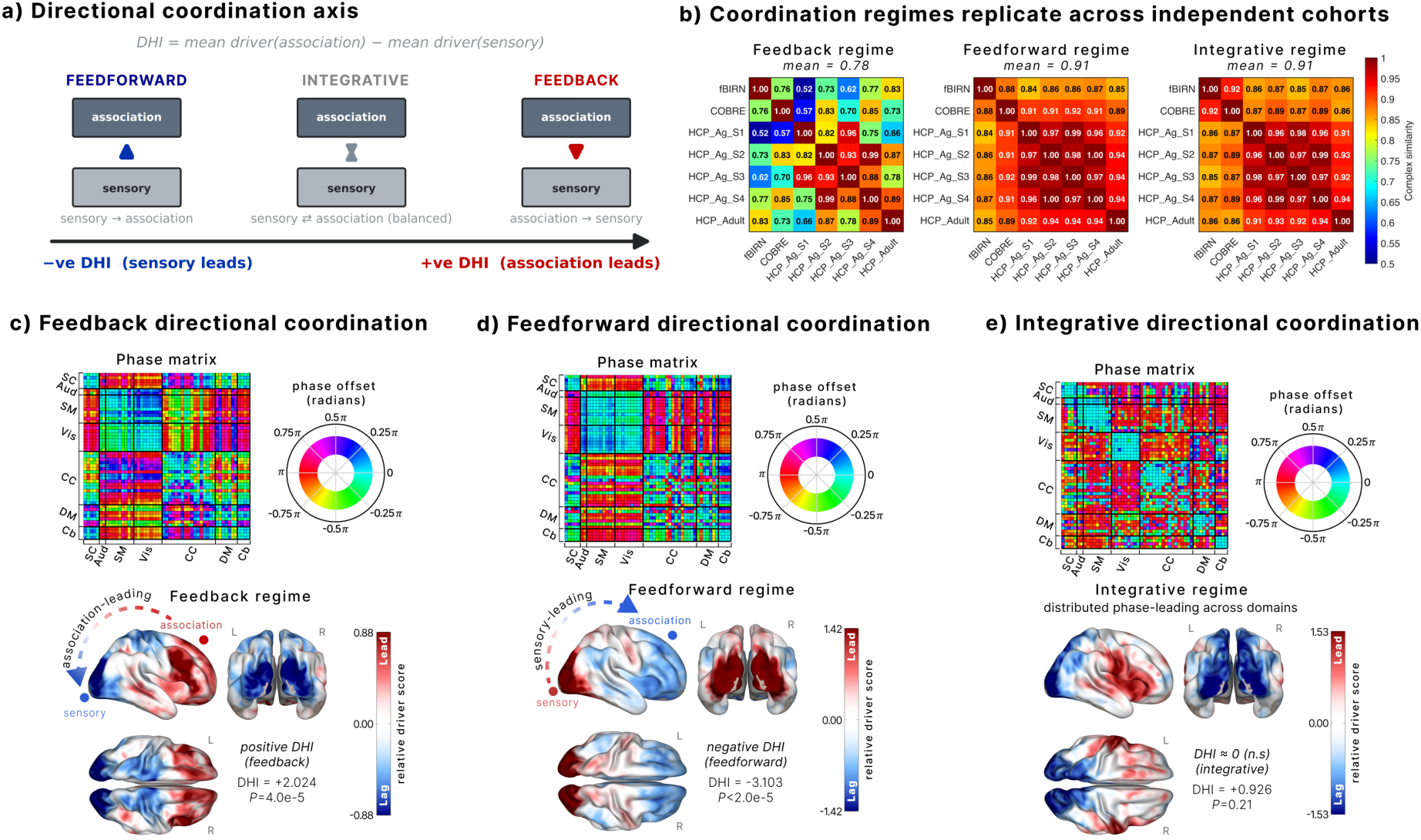
Directional coordination forms three reproducible regimes. **(a)** Directional coordination axis. The directional hierarchy index (DHI) is the mean driver score of association minus sensory systems: positive indicates association-leading (top-down, feedback), negative indicates sensory-leading (bottom-up, feedforward), and near zero indicates balanced exchange (integrative). **(b)** Regimes replicate across independent cohorts. Pairwise complex-correlation matrices of regime centroids across seven datasets from four independent cohorts (fBIRN, COBRE, HCP-Aging sessions 1–4, HCP-Adult). Mean similarity was 0.78 for feedback and 0.91 for both feedforward and integrative, reproducing across acquisition protocols, scanner sites, and populations. **(c to e)** Three directional coordination regimes from complex-valued phase synchrony (CVPS) in the discovery cohort (fBIRN, n = 311). Per regime: phase matrix across functional domains (left; subcortical (SC), auditory (Aud), sensorimotor (SM), visual (Vis), cognitive control (CC), default mode (DM), cerebellum (Cb)), phase-offset color wheel (center), and cortical projection of relative driver scores (right; red leading, blue lagging). **(c)** Feedback: association leads sensory (DHI = +2.024, p = 4.0 × 10⁻⁵). **(d)** Feedforward: sensory leads association (DHI = −3.103, p < 2 × 10⁻⁵). **(e)** Integrative: balanced phase-leading with no dominant hierarchy (DHI = +0.926, p = 0.21, n.s.). P-values are from 50,000 random network label permutations.

The same three regimes recurred across independent cohorts spanning different acquisition protocols, scanner sites, and populations, comprising COBRE, four sessions of HCP-Aging, and HCP-Adult. Complex correlation between regime representations gave a mean cross-dataset similarity of 0.78 for the feedback regime and 0.91 for both the feedforward and integrative regimes (Fig. 1b).

Each regime carried a distinct configuration of phase offsets across canonical functional systems. In the feedback regime, association regions systematically led sensory systems, yielding a positive index (DHI = +2.024, p = 4.0 × 10⁻⁵; Fig. 1c). The feedforward regime showed the opposite organization, with sensory regions leading association systems (DHI = −3.103, p < 2 × 10⁻⁵; Fig. 1d). In the integrative regime, no domain consistently led, and the index did not depart from zero (DHI = +0.926, p = 0.21; Fig. 1e). The axis thus spans two directional poles and a balanced integrative regime, each reproducible across cohorts.

### 3.2. Directional coordination is selectively reorganized in schizophrenia and replicates in an independent cohort

Beyond discrete regimes, directional coordination occupies a continuous persistence landscape in which basin depth integrates each regime’s self-persistence with its long-run equilibrium weighting from the Markov structure of regime transitions, yielding well-separated feedback, feedforward, and integrative basins in both patients and controls (Fig. 2a). In schizophrenia (fBIRN; 151 patients, 160 controls), this landscape did not collapse uniformly but reorganized selectively: the integrative basin deepened while the feedback basin flattened, with the feedforward basin comparatively spared (Fig. 2b; p < 0.001).

**Fig. 2:**
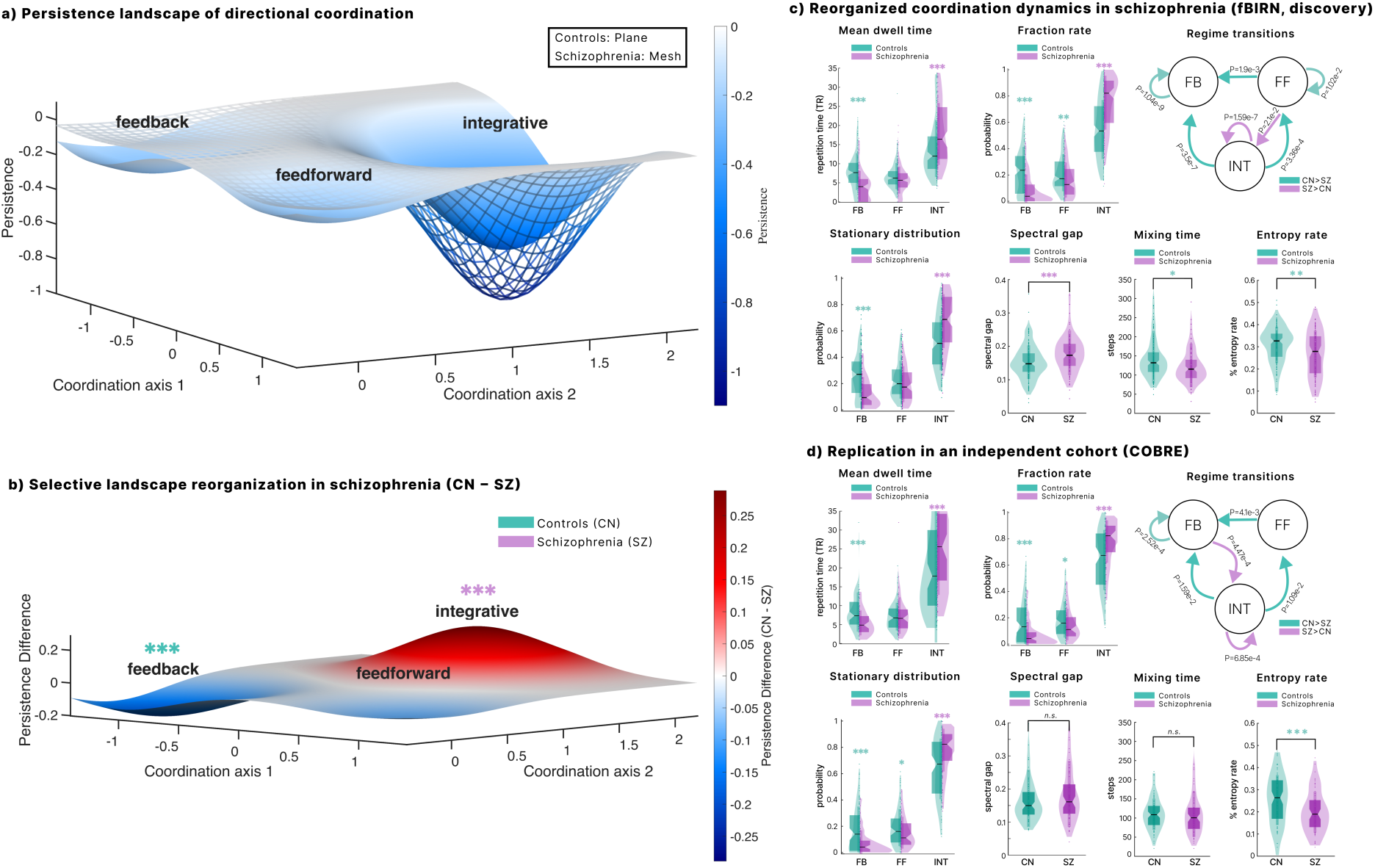
Directional coordination is selectively reorganized in schizophrenia and replicates in an independent cohort. **(a)** Persistence landscape of directional coordination. Low-dimensional projection of coordination dynamics, with surface depth indexing persistence (basin depth reflects regime stability and preferential fraction rate). Controls are shown as a smooth surface and schizophrenia as an overlaid mesh, revealing separable feedback, feedforward, and integrative basins in both groups (fBIRN; 160 controls, 151 individuals with schizophrenia). **(b)** Selective landscape reorganization in schizophrenia (controls minus schizophrenia). Controls show deeper persistence in the feedback basin, whereas schizophrenia shows deeper persistence in the integrative basin (red), a selective redistribution of stability, not a uniform collapse (*** p < 0.001). **(c)** Group differences in directional coordination dynamics in the discovery cohort (fBIRN), from generalized linear models controlling for age, sex, scanner site, and head motion, FDR-corrected. Mean dwell time, fraction rate, and the stationary distribution indicate reduced feedback and increased integrative engagement in schizophrenia; the regime-transition diagram shows altered flow between regimes (teal: controls greater; purple: schizophrenia greater). Global dynamical properties shift consistently: schizophrenia shows a higher spectral gap and shorter mixing time (faster convergence to equilibrium) and a lower entropy rate (less diverse trajectories). **(d)** Replication in an independent cohort (COBRE; 89 controls, 68 individuals with schizophrenia), analyzed identically except that scanner site is omitted as COBRE is single-site. Feedback erosion, integrative deepening, reduced entropy rate, and the rerouting of transitions toward integrative coordination all reproduce; spectral gap and mixing time do not differ significantly (n.s.), though in the same direction as the discovery cohort. Asterisks indicate FDR-corrected significance in (b), (c), and (d) (* p < 0.05, ** p < 0.01, *** p < 0.001). FB, feedback; FF, feedforward; INT, integrative.

Metric-level comparisons revealed the same result (Fig. 2c). Relative to controls, patients spent less time in feedback and more in integrative coordination across mean dwell time, fraction rate, and the long-run stationary distribution, and their transition structure rerouted flow away from the directional regimes toward integrative coordination. Global dynamical properties shifted in the same direction. Schizophrenia showed a higher spectral gap and shorter mixing time, indicating faster convergence to equilibrium, together with a lower entropy rate, indicating less diverse temporal trajectories. The dynamics in schizophrenia settled rapidly into the integrative regime and explored a narrower portion of the coordination repertoire.

To test whether this reorganization is a reproducible feature of the disorder and not a property of one cohort, we repeated the full analysis in an independent clinical sample (COBRE; 68 patients, 89 controls; Fig. 2d). The selective pattern reproduced: feedback dwell time, fraction rate, and stationary distribution were reduced, integrative dwell time, fraction rate, and stationary distribution were increased, entropy rate was reduced, and transitions rerouted toward integrative coordination, all at FDR-corrected significance. Spectral gap and mixing time did not differ significantly in this smaller sample, though the direction was consistent with that of the discovery cohort. Of the seventeen effects significant in the discovery cohort, thirteen replicated at FDR-corrected significance in COBRE. The reorganization of directional coordination in schizophrenia is therefore selective, convergent across dynamical metrics, and reproducible across independent cohorts.

### 3.3. Directional coordination dynamics track positive symptoms, and their relationship to cognition reverses in schizophrenia

The reorganization was behaviorally and clinically meaningful, and its relationship to cognition was not merely weakened in schizophrenia but reversed (Fig. 3). In controls, more frequent transitions from feedback toward integrative coordination tracked lower attention vigilance and lower composite cognition, while faster convergence to equilibrium, reflected in a higher spectral gap and shorter mixing time, tracked better verbal learning (Fig. 3a). In schizophrenia, longer feedback dwell time tracked better attention vigilance, and the convergence relationships took the opposite sign. The same faster convergence that accompanied better verbal learning in controls tracked worse attention vigilance and worse composite cognition in patients (Fig. 3b). The normative link between coordination dynamics and cognition is therefore not simply diminished in schizophrenia but takes an opposite-signed form.

**Fig. 3:**
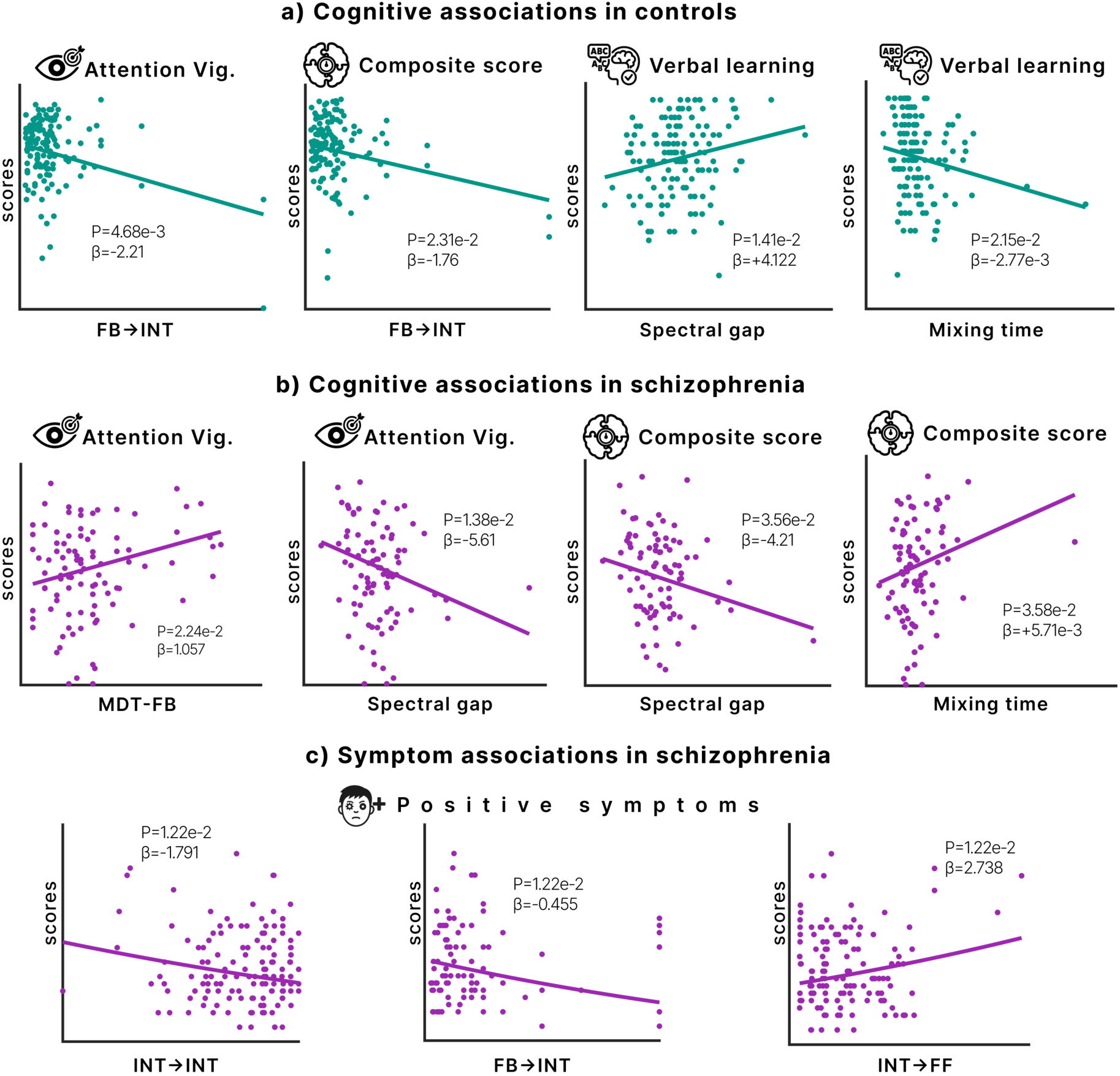
Coordination dynamics track positive symptoms and relate to cognition in opposite directions in schizophrenia. Within-group associations linking directional coordination dynamics to cognitive and clinical measures (generalized linear models controlling for age, sex, scanner site, and head motion; FDR-corrected). Each scatter shows the regression fit with its regression coefficient and FDR-adjusted p-value. **a)** Cognitive associations in controls (n = 160). More frequent feedback-to-integrative transitions track lower attention vigilance and lower composite cognition, while faster convergence to equilibrium (higher spectral gap, shorter mixing time) tracks better verbal learning. **b)** Cognitive associations in schizophrenia (n = 151). Longer feedback dwell time tracks better attention vigilance, and, in contrast to controls, faster convergence (higher spectral gap, shorter mixing time) tracks worse attention vigilance and worse composite cognition. **c)** Symptom associations in schizophrenia. Greater persistence of integrative coordination and more frequent feedback-to-integrative transitions track lower positive symptom severity, whereas more frequent integrative-to-feedforward transitions track higher positive symptom severity. FB, feedback; FF, feedforward; INT, integrative; MDT, mean dwell time.

Positive symptom severity was tied to transition structure (Fig. 3c). Greater persistence of integrative coordination and more frequent feedback-to-integrative transitions tracked lower positive symptoms, whereas more frequent transitions from integrative toward feedforward coordination tracked higher positive symptoms. Sustained access to integrative coordination thus appears relatively adaptive, while slipping from it toward feedforward dynamics indexes symptom expression. Results were unchanged when chlorpromazine-equivalent dose was additionally included as a covariate within patients (Supplementary Table ST. 1).

### 3.4. GABAergic sedation and motor imagery shift directional coordination toward opposite poles within the same subjects

The group differences above presuppose that the directional axis is a controllable property of brain dynamics and not an arbitrary partition of a clustering solution. We therefore tested whether the axis can be moved experimentally in a cohort not used in any analysis above: 26 healthy volunteers (OpenNeuro ds006623), scanned under two within-subject manipulations and parcellated with an anatomically defined atlas as opposed to the ICA-derived components from earlier analyses (Fig. 4a). The same three regimes re-emerged with their directional signatures intact (Fig. 4b, c). Sensory networks led in the feedforward regime (DHI = −44.7, p < 10⁻⁴), association networks led in the feedback regime (DHI = +36.78, p < 10⁻⁴), and phase-leading was balanced in the integrative regime (DHI = +0.66, p = 0.83). Because this parcellation is atlas-based, the recurrence of the regimes establishes them as a reproducible feature of cortical organization and not a parcellation-specific artifact.

**Fig. 4:**
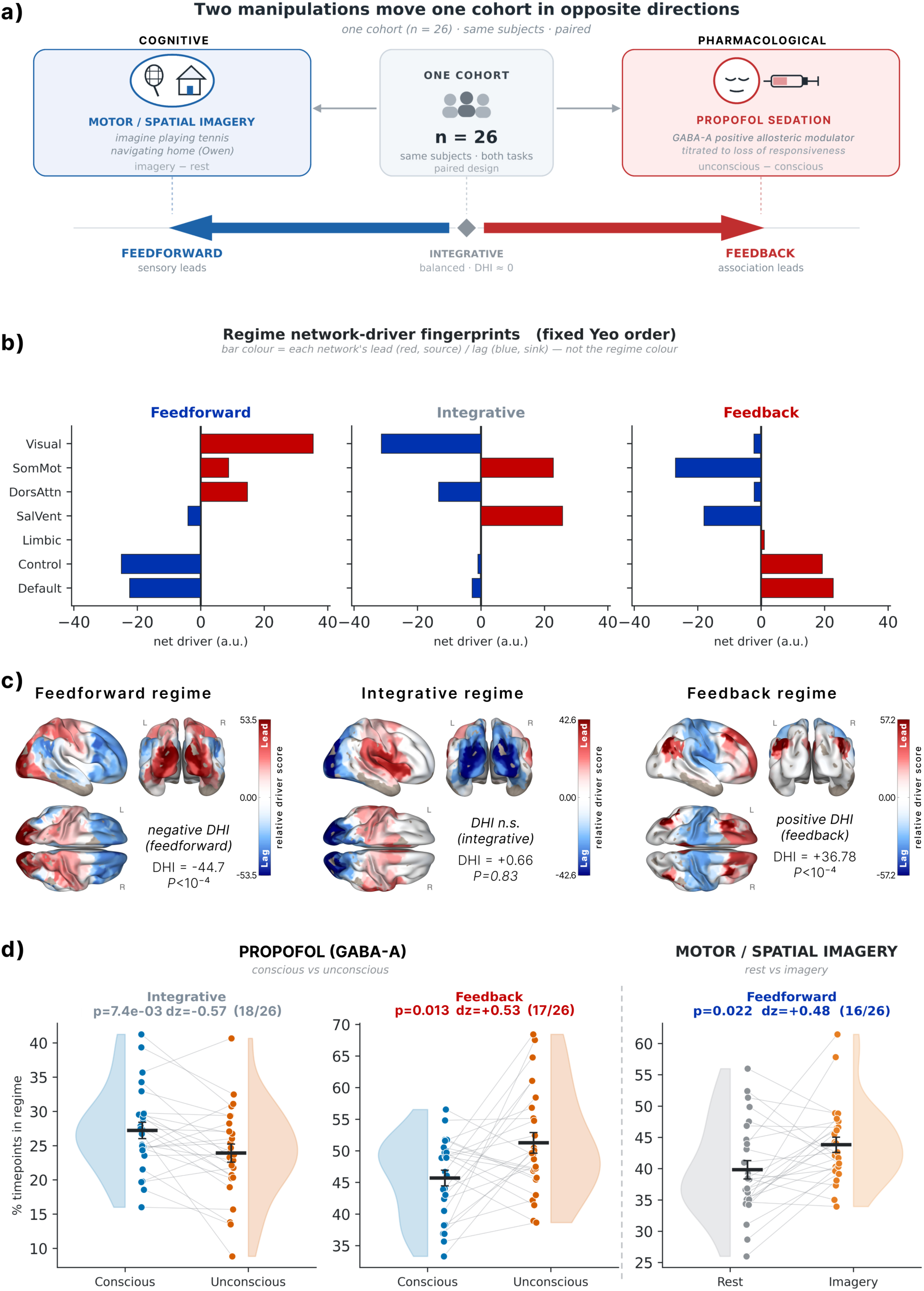
GABAergic sedation and motor imagery shift directional coordination toward opposite poles in the same subjects. **(a)** Design. One cohort (n = 26, OpenNeuro ds006623, paired) underwent two within-subject manipulations predicted a priori to move coordination toward opposite poles: motor and spatial imagery (imagery vs rest) toward feedforward, and propofol, a GABA-A positive allosteric modulator titrated to loss of responsiveness (unconscious vs conscious), toward feedback. **(b)** Net directional drive averaged within the seven Yeo networks for each regime (Glasser-360 atlas); bar color denotes lead (red) or lag (blue), not regime. **(c)** Cortical relative driver scores (red leading, blue lagging) with the directional hierarchy index (DHI) per regime: feedforward −44.7 (p < 10⁻⁴), integrative +0.66 (p = 0.83, n.s.), feedback +36.78 (p < 10⁻⁴). Recovery of the same three regimes under an anatomical atlas, not the ICA-derived components used at discovery, constitutes replication across parcellations; DHI is in arbitrary units and not comparable to Fig. 1. **(d)** Within-subject change in regime fraction rate (lines connect subjects). Paired Cohen’s dz, FDR-corrected, with the number shifting in the predicted direction in parentheses; dz is manipulation minus baseline, so positive indicates increased fraction rate. Sedation increased feedback (dz = +0.53, p = 0.013, 17/26) and decreased integrative fraction rate (dz = −0.57, p = 7.4 × 10⁻³, 18/26) without increasing feedforward (p = 0.17, n.s.); imagery increased feedforward (dz = +0.48, p = 0.022, 16/26) without changing integrative (p = 0.09) or feedback (p = 0.14). Each manipulation moved its predicted regime and spared the opposite pole.

Each manipulation was predicted a priori to move coordination toward an opposite pole of the axis (Fig. 4a). Propofol, a positive allosteric modulator at GABA-A receptors^42^ titrated to loss of responsiveness, was predicted to shift coordination toward the feedback pole, and motor and spatial imagery toward the feedforward pole. Because both manipulations were applied to the same subjects, any shift must be intrinsic to those subjects rather than a difference between groups or scanners.

Both predictions held, and each manipulation moved only its predicted pole (Fig. 4d). Sedation increased feedback fraction rate (paired Cohen’s dz = +0.53, p = 0.013, 17 of 26 subjects) and decreased integrative fraction rate (dz = −0.57, p = 7.4 × 10⁻³, 18 of 26), without increasing feedforward fraction rate (p = 0.17). Imagery increased feedforward fraction rate (dz = +0.48, p = 0.022, 16 of 26), without changing the feedback (p = 0.14) or integrative (p = 0.09) regimes. A pharmacological and a cognitive manipulation therefore produced a within-subject double dissociation, each shifting coordination toward its predicted pole while sparing the opposite one.

### 3.5. Directional coordination is hierarchical, not a fixed vascular lag, and distinct from neighboring principles

We next asked whether the directional ordering within each regime reflects the cortical hierarchy as defined independently of our analysis. We correlated each region’s net directional driver with its position on the Margulies principal gradient, a static spatial axis computed from resting-state connectivity in a separate sample of 207 unrelated HCP adults distributed with the ENIGMA toolbox (Fig. 5a). Neither our lead-lag measures nor our sensory-association grouping enters its construction. The alignment reversed by regime exactly as a directional hierarchy predicts. Association regions led and correlated positively with the gradient in the feedback regime (Spearman rho = +0.76, p = 0.002); sensory regions led association regions. The correlation reversed in the feedforward regime (rho = −0.64, p < 10⁻³), and the integrative regime showed no alignment (rho = −0.18, p = 0.486). Both directional poles exceeded a spatial-autocorrelation-preserving null, and the alignment was stronger under the anatomical parcellation of the manipulation cohort (rho = +0.71 and −0.83; Supplementary Fig. SF. 3). The gradient was constructed without our data, and our regimes were defined without the gradient, so this correspondence is discovered, not built in.

**Fig. 5:**
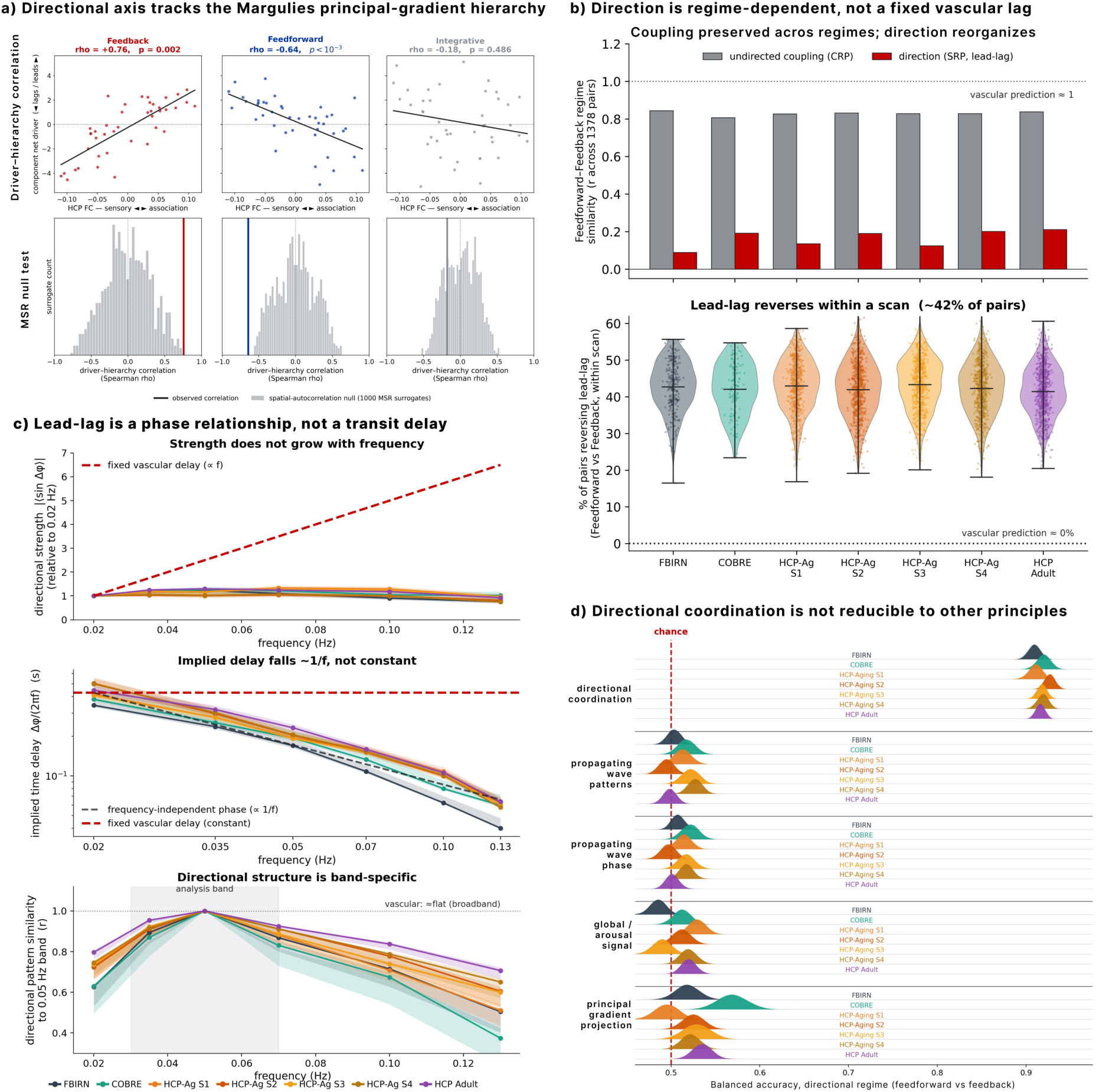
Directional coordination is hierarchical, not a fixed vascular lag, and distinct from neighboring principles. **(a)** Per-region net driver in each regime against the Margulies principal gradient, an external hierarchy computed from independent HCP connectivity and blind to our lead-lag and grouping (fBIRN, cortical NeuroMark components). The alignment reverses by regime exactly as a directional hierarchy predicts: feedback drivers correlate positively (association leads), feedforward negatively (sensory leads), and integrative drivers not at all. Both directional poles lie outside a spatial-autocorrelation-preserving null, and the integrative regime within it (1,000 Moran surrogates, gray; bottom); therefore, the alignment is not an artifact of spatial smoothness. **(b)** The two most directionally opposite regimes contrasted across 1,378 region pairs per subject. Where a fixed vascular delay would give identical direction in every regime (similarity near 1, near 0% reversals), coupling was instead preserved (CRP, gray; r = 0.81 to 0.85) while direction reorganized (SRP, red; r = 0.1 to 0.2), with roughly 42% of pairs reversing lead-lag sign within a single scan, so that the same pair both leads and lags in one recording. This implies that directional coordination is not imposed by fixed vascular timing but reconfigures within a scan. **(c)** A fixed conduction delay makes three predictions (red dashed), and all three fail: directional strength should rise with frequency but stays flat from 0.02 to 0.13 Hz (top); the implied delay should be constant but falls as roughly 1/f (middle); and the directional pattern should hold broadband but instead peaks in the 0.03 to 0.07 Hz analysis band and decorrelates outside it (bottom), which implies that directional coordination is not a fixed transit delay but a band-limited phase relationship. Solid lines are cohort means with 95% confidence intervals. **(d)** At each time point, the active regime (feedforward or feedback) was decoded from five feature sets, with no subject in both training and test folds (balanced accuracy; chance = 0.5). The lead-lag features themselves reach approximately 0.92 in every cohort, a ceiling since the regimes are defined in that space. Propagating-wave patterns, propagating-wave phase, the global/arousal signal, and the principal gradient projection all sit at chance, which implies that the feedforward-versus-feedback distinction is not contained in these constructs.

A directed lead-lag could in principle reflect a fixed hemodynamic transit delay instead of coordination. A fixed delay is static, predicting identical direction in every regime and no within-scan reversals, and neither held (Fig. 5b). Across the two most directionally opposite regimes, coupling was preserved over the 1,378 region pairs (CRP, r = 0.81 to 0.85) while direction reorganized almost completely (SRP, r = 0.1 to 0.2), and roughly 42% of pairs reversed their lead-lag sign within a single scan. The same pair both leads and lags within one recording, which no static delay can produce. A fixed delay also predicts directional strength rising linearly with frequency, a constant implied delay, and a frequency-flat pattern, and the data violated all three (Fig. 5c). Strength stayed flat across 0.02 to 0.13 Hz, the implied delay fell as roughly 1/f, and the directional pattern peaked in the 0.03 to 0.07 Hz analysis band and decorrelated elsewhere. Lead-lag is thus a band-limited, regime-dependent phase relationship and not a fixed conduction delay. Both control analyses held across all seven datasets.

Finally, we asked whether the directional distinction simply re-describes other known organizing principles (Fig. 5d). Using subject-grouped cross-validation, we decoded the active regime (feedforward versus feedback) from each feature set, scoring performance as balanced accuracy, the average accuracy across the two regimes, so that unequal fraction rate cannot inflate it (chance = 0.5). The lead-lag features themselves decoded the regime at approximately 0.92 in every cohort, confirming the partition is real and learnable. None of the four neighboring constructs, propagating-wave patterns, propagating-wave phase, the global arousal signal, or the principal gradient projection, decoded it above chance in any dataset. The same gradient that predicts which regions lead within a regime (Fig. 5a) therefore carries no information about which regime is active (Fig. 5d). The static cortical hierarchy specifies where coordination flows, and only directional coordination specifies when.

## 4. DISCUSSION

We found that ongoing brain activity is organized along a directional coordination axis that aligns region by region with an externally defined cortical hierarchy. Three regimes recur along this axis across independent cohorts, one in which association cortex leads sensory systems (feedback), one in which sensory systems lead (feedforward), and one of balanced bidirectional exchange (integrative). In schizophrenia, the axis reorganized selectively. Feedback coordination lost persistence, integrative coordination gained it, and the dynamics converged faster onto a narrower repertoire, a pattern that reproduced in an independent clinical cohort and tracked positive symptoms and cognition. Within the same healthy subjects, a GABA-A receptor challenge and a cognitive task moved the axis in opposite directions, establishing it as a controllable property of brain dynamics. Directional coordination is invisible to conventional connectivity measures, which discard the sign of lead-lag relationships, and it was not recoverable from propagating waves, arousal, or the principal gradient.

Schizophrenia has long been framed as a disorder of hierarchical dysconnection, in which aberrant precision-weighting of prediction errors destabilizes the top-down signals that normally constrain sensory representations^1,2,5^. Our results give this account a dynamical form. The feedback regime, in which association cortex leads sensory systems, corresponds to the descending pathway of hierarchical processing, and the feedforward regime to the ascending one^3,4,15,16^. Dysconnection in schizophrenia has chiefly been characterized as altered strength of coupling between regions^27,59^. Our findings locate a deficit in a dimension those measures do not capture. The persistence of the association-leading pole erodes while the feedforward pole is comparatively spared, a selective reorganization of directional structure consistent with evidence for compressed hierarchical separation between sensory and association systems in the disorder^23,24^. That greater integrative fraction rate is common at rest yet elevated in patients is resolved by the distinction between a visited regime and a terminal one^60^. Balanced exchange is adaptive as one of three regimes the system moves through, and maladaptive when the dynamics collapse into it. The faster convergence and reduced entropy rate we observed indicate such a collapse, a reduced capacity to sustain excursions away from equilibrium, deepening prior reports of reduced connectivity dynamism in schizophrenia^27–29^ by locating the deficit at the level of direction. Model-based evidence points the same way, with weakened top-down and strengthened bottom-up effective connectivity reported at rest in small prefrontal-sensory circuits^12^.

The sedation result requires care, because it appears at first to contradict both our schizophrenia findings and a large anesthesia literature. Propofol shifted occupancy toward the feedback pole and away from integration (Fig. 4d), the mirror image of the schizophrenia profile (Fig. 2b), yet decades of work show that anesthesia preferentially disrupts top-down connectivity^61,62^. The resolution lies in what the feedback pole measures. The regime is defined by geometry, association cortex leading sensory cortex, and more than one process can produce that geometry at the hemodynamic timescale. Deep sedation is dominated by an anteroposterior slow oscillation that originates in the frontal cortex and propagates backward^63,64^, which is association-leading geometry without the flexible top-down communication that anesthesia is known to abolish. The propofol shift therefore marks entry into a stereotyped, anteriorly driven slow-wave mode, and the concurrent loss of integrative coordination matches the collapse of flexible coordination repertoires that characterizes unconsciousness^65,66^. Viewed this way, sedation and schizophrenia occupy a common axis and fail in opposite directions. Each collapses the coordination repertoire toward a rigid attractor: sedation toward the slow-wave feedback pole with loss of integration, schizophrenia toward the integrative pole with loss of feedback. The shared signature is reduced flexibility, and the directional framework specifies which mode each condition forfeits.

The relationship between coordination dynamics and cognition was not merely weakened in patients but inverted. In controls, faster convergence to equilibrium accompanied better verbal learning, consistent with evidence that the structure of resting information flow predicts cognitive ability^34,67^. In schizophrenia, the same convergence tracked worse attention and worse composite cognition. A sign inversion is informative in a way that an attenuation is not, because reduced coupling alone cannot change the direction of an association. It suggests that rapid settling serves cognition when the settled dynamics retain directional structure, and costs cognition when settling means collapse into an undirected mode, mirroring evidence that lost dynamical flexibility underlies cognitive impairment in the disorder^68,69^. Symptom expression showed the complementary pattern. Transitions into integrative coordination tracked lower positive symptoms while transitions toward feedforward coordination tracked higher ones, consistent with accounts in which positive symptoms arise when ascending signals go uncontextualized by top-down prediction^5,6^. Together, these associations point to potential clinical uses. A measure whose cognitive coupling differs in sign between patients and controls is a potential stratification readout, symptom-linked transition structure offers a monitoring target, and the axis’ sensitivity to a GABA-A challenge suggests a possible target-engagement marker for interventions aimed at hierarchical dynamics, each contingent on longitudinal validation.

Some organizing principles adjoin the one described here, and none contains it. Gradients of intrinsic timescale order regions by how long their activity persists^19,20^, propagating waves describe activity traversing the hierarchy on average^21,22^, arousal fluctuations impose global coordination^43^, and the principal gradient arranges regions by the similarity of their connectivity^18^. Each captures a time-averaged or undirected property, and our decoding analyses showed that none recovers which directional regime is active at a given moment. Earlier lag analyses of resting fMRI identified reproducible temporal orderings^38^ but averaged them over entire scans, which collapses regimes of opposite direction onto one another. Phase-based dynamic connectivity resolved coordination in time but discarded the sign of the phase relationship by convention^31–33,35^. Directed models such as Granger causality and dynamic causal modeling estimate influence within small networks under strong model assumptions^11,40^. Convergent work on temporal irreversibility indicates that large-scale brain dynamics possess an arrow of time^70^. A thermodynamic account has quantified hierarchical asymmetry in schizophrenia on this footing, but as a single scalar that does not say which levels lead^71^. Directional coordination specifies, region by region and moment by moment, which way that arrow points.

Several limitations bound these conclusions. Although the CVPS method applies to any time series, we applied it here to fMRI, whose hemodynamic timescale reflects the slow envelope of neural activity and not the millisecond signaling of cortical transmission^72^. Our controls exclude a fixed vascular delay, but they do not establish a neural origin, and state-dependent hemodynamic variation remains a possible contributor, particularly under propofol, which alters vascular reactivity alongside neural dynamics. However, against a purely vascular account, the propofol shift was regime-selective, and imagery moved the same axis with no pharmacological action. Applying the same framework to electrophysiology, where gamma-band feedforward and alpha- and beta-band feedback channels are established^17^, would address both concerns. The cross-sectional design, in chronic medicated samples, cannot determine whether feedback erosion precedes illness onset. Longitudinal designs spanning the clinical high-risk to first-episode transition, and extension to other disorders of hierarchical dynamics, are natural next steps.

Large-scale brain dynamics have been characterized chiefly by which states recur and how often. Direction, the question of which regions lead and which follow, has been treated as a property to be averaged away or implicitly ignored. Treating it as signal revealed a dimension of coordination that replicates across cohorts, aligns with the cortical hierarchy, responds to pharmacological and cognitive manipulation, and is selectively disrupted in schizophrenia in proportion to symptoms and cognition.

## 6. AUTHOR CONTRIBUTION

**Sir-Lord Wiafe:** Conceptualization, Formal analysis, Methodology, Visualization, Writing – original draft. **Najme Soleimani:** Validation, Writing – review & editing. **Zening Fu:** Data analysis, Validation, Writing – review & editing. **Robyn Miller:** Conceptualization, Validation, Supervision, Writing – review & editing. **Vince Calhoun:** Conceptualization, Funding acquisition, Validation, Methodology, Resources, Supervision, Writing – review & editing.

## 7. DATA & CODE AVAILABILITY STATEMENT

The replication and validation datasets are publicly available: COBRE via the Collaborative Informatics and Neuroimaging Suite, HCP-Aging and HCP-Adult via the Human Connectome Project, and the task and anesthesia cohort via OpenNeuro (ds006623). fBIRN data are available through the fBIRN data repository. Individual-subject fBIRN derivatives cannot be redistributed by the authors, as no data-reuse agreement was signed at the original acquisition. Analysis code implementing CVPS, regime identification, the DHI, and the control analyses is available on <u>GitHub</u>.

## 8. ACKNOWLEDGMENT

This work was supported by the National Institutes of Health (NIH) grant (R01MH123610) and the National Science Foundation (NSF) grant #2112455.

