## supplementary material for "Directional coordination along the cortical hierarchy is selectively reorganized in schizophrenia"

### 1. Supplementary fMRI data

#### 1.1. fBIRN discovery cohort

We analyzed resting-state fMRI from Phase III of the Function Biomedical Informatics Research Network (fBIRN) consortium[1], comprising 160 healthy controls (37.0 ± 10.9 years, 45 F/115 M) and 151 individuals with schizophrenia (38.8 ± 11.6 years, 36 F/115 M) scanned at seven sites. Data were acquired with a repetition time of 2 s. Preprocessing discarded the initial 10 volumes and comprised slice-timing correction, rigid-body realignment, spatial normalization to MNI space, and smoothing with a 6 mm full-width at half full-width-at-half-maximum Gaussian kernel. Spatially independent components were extracted using the NeuroMark pipeline with the neuromark_fMRI_1.0 template[2] (available in GIFT at http://trendscenter.org/software/gift), yielding 53 intrinsic connectivity networks (ICNs) per subject, grouped into subcortical, auditory, sensorimotor, visual, cognitive control, default mode, and cerebellar domains. ICN time courses were z-scored before CVPS analysis. Symptom severity was assessed with the Positive and Negative Syndrome Scale (PANSS) and cognition with the Computerized Multiphasic Interactive Neurocognitive System (CMINDS).

#### 1.2. COBRE cohort

We analyzed resting-state fMRI from the Center for Biomedical Research Excellence (COBRE) dataset[3] as the independent clinical replication cohort for the schizophrenia analyses (Fig. 2d) and as a healthy replication cohort for regime identification. The present analysis comprised 89 healthy controls (38.1 ± 11.7 years, 64 M) and 68 individuals with schizophrenia (37.8 ± 14.4 years, 57 M), scanned at a single site. Resting-state fMRI data were acquired during an approximately 5-minute resting-state scan using a repetition time of 2 s and echo time of 29 ms, yielding 150 time points. Volumes underwent the same preprocessing and network-level extraction framework used in the discovery analysis. Spatially independent components were extracted using the NeuroMark pipeline with the neuromark_fMRI_1.0 template, yielding 53 intrinsic connectivity networks (ICNs) for each subject. The resulting ICN time courses were z-scored and submitted to the same complex Gabor-wavelet phase-synchrony analysis used for fBIRN. PANSS scores were available for all patients.

#### 1.3. HCP aging cohort

We analyzed resting-state fMRI from the Human Connectome Project Aging dataset[4] as an independent lifespan replication cohort. The HCP Aging dataset includes multimodal neuroimaging and behavioral data from healthy adults spanning later adulthood, with participants aged 36 to 100+ years. The present analysis comprised 718 participants. Resting-state fMRI data were acquired using HCP-style multiband echo-planar imaging in four runs of approximately 6.5 minutes each, yielding approximately 26 minutes of resting-state data at a TR = 800 ms. Each run included 488 frames, with alternating phase-encoding directions across runs. Also, each run was analyzed as a separate dataset (sessions S1 to S4 in the main text), providing four within-cohort replications. Spatially independent components were extracted using the NeuroMark pipeline with the neuromark_fMRI_1.0 template, yielding 53 intrinsic connectivity networks (ICNs) for each subject. The resulting ICN time courses were z-scored and submitted to the same complex Gabor-wavelet phase-synchrony analysis used for fBIRN.

##### *1.4. HCP adult cohort*

We analyzed resting-state fMRI from the Human Connectome Project Young Adult dataset[5] as an independent healthy adult replication cohort. The HCP Young Adult dataset includes high-quality multimodal neuroimaging and behavioral data from healthy adults aged 22 to 35 years. The present analysis comprised 827 participants. Resting-state fMRI data were acquired across four runs of approximately 15 minutes each, with two runs collected in one session and two runs collected in a second session. Participants viewed a projected fixation cross during rest. Oblique axial acquisitions alternated between right-to-left and left-to-right phase encoding directions. Images were acquired using gradient-echo EPI with TR = 720 ms, TE = 33.1 ms, flip angle = 52°, 2.0 mm isotropic voxels, multiband factor = 8, and 1200 frames per run. Spatially independent components were extracted using the NeuroMark pipeline with the neuromark_fMRI_1.0 template, yielding 53 intrinsic connectivity networks (ICNs) for each subject. The resulting ICN time courses were z-scored and submitted to the same complex Gabor-wavelet phase-synchrony analysis used for fBIRN.

##### *1.5. Task and anesthesia cohort (OpenNeuro ds006623)*

Twenty-six healthy volunteers (ages 19–34; 13 women) were scanned on a 3 T Philips scanner with a 32-channel head coil[6]. Functional images used a gradient-echo EPI sequence with 28 slices, TR/TE = 800/25 ms, multiband factor 4, 4 mm slice thickness, 3.4 × 3.4 mm in-plane resolution, FOV = 220 mm, flip angle = 76°, and a 64 × 64 matrix; six participants used a modified protocol (21 slices, multiband factor 3, 6 mm slice thickness, TR/TE unchanged). A T1-weighted anatomical image was acquired with 170 sagittal slices, 1.0 mm thickness, TR = 8.1 s, TE = 3.7 ms, flip angle = 8°, FOV = 240 mm, 256 × 256 matrix. Sedation used propofol titrated to target effect-site concentrations (initial 0.4 or 1.0 µg/ml; final 2.4 µg/ml or one 0.4 µg/ml increment above loss of responsiveness). The session comprised two 10-minute resting-state runs and task runs spanning a 15-minute awake baseline, a 30-minute propofol-sedation scan, a 30-minute post-infusion scan, and a 15-minute recovery baseline; the imagery block design alternated 15 s of tennis or spatial-navigation imagery, 10 s of hand-squeeze imagery, and 15 s of rest[7]. We analyzed the preprocessed time series distributed with the dataset (fMRIPrep, normalized to MNI152NLin6Asym; XCP-D denoising), parcellated with the Glasser-360 atlas (HCP-MMP1.0)[8], and mapped regions to the seven Yeo networks using the community labels provided with the segmentation[9].

##### *1.6. External gradient reference sample*

The principal functional connectivity gradient[10] was computed from group-average resting-state functional connectivity of 207 unrelated healthy adults from the Human Connectome Project, distributed with the ENIGMA toolbox[11]. These participants are distinct from all cohorts analyzed above, and neither our lead-lag measures nor our system grouping enters the gradient's construction.

### 2. Complex-valued phase synchrony mathematics

The complex Gabor wavelet defined in the main text (Equation 1) was truncated and normalized before convolution. To reduce computation while retaining $99.7\%$ of the Gaussian energy, the kernel is truncated to $t \in [-3\sigma_t, +3\sigma_t]$[12], sampled on the TR grid $(t = k\Delta t, k \in \mathbb{Z})$. The analytic signal is

$$z(t) = (x * g)(t) = \sum_k x(k\Delta t) g(t - k\Delta t) \Delta t \qquad \text{(SE. 1)}$$

with $g(t)$ normalized to unit energy to avoid edge bias. The instantaneous phase and amplitude follow as:

$$\varphi(t) = \arg[z(t)], \quad A(t) = |z(t)| \qquad \text{(SE. 2)}$$

The phase $\varphi(t)$ so obtained is the quantity entering the CVPS definition in the main text (Equation 2). Relative to a Hilbert band-pass pipeline, the adaptive Gabor kernel affords sharper time-frequency localization for the non-stationary BOLD spectrum while preserving the full complex phase necessary for synchrony analysis[13], as demonstrated in the simulations below (Fig. SF. 1e).

CVPS can be understood within the dynamic co-modulation (DyCoM) framework, which expresses time-varying interactions through a small set of fundamental signal processing operations[14]. In this framework, CVPS is a phase-based instantiation in which the representation operator is the Gabor wavelet transform, and the interaction operator is the complex phase difference, without any instantiation of the temporal or normalization operators.

### 3. Simulation studies

#### *3.1.* Necessity of preserving the full complex angle (Fig. SF. 1a)

To show that cosine-based synchrony cannot separate configurations that differ only in lead-lag direction, we generated two synthetic datasets of phase-coupled sinusoids. Each set comprised N = 100 signal pairs, created as 0.03 Hz sine waves sampled at 1.5 Hz for 666 seconds (1,000 samples). The second signal of every pair was phase-shifted by +60° (Group A) or –60° (Group B), with Gaussian jitter ($\sigma = 10°$) added on each trial to mimic variability. Instantaneous phases were extracted with the Hilbert transform, yielding a stationary pairwise phase difference $\Delta\varphi$. For every pair, we computed the classical cosine synchrony $cos(\Delta\varphi)$ and the direction-preserving mean vector angle $arg(e^{j\Delta\varphi})$. Group differences were assessed with a two-sample t-test for the cosine values and Watson's $U^2$ circular-statistics test for the mean-vector angles. The two groups are indistinguishable in the cosine ($P = 0.74$) and cleanly separable in the full angle ($P \approx 0$), establishing that opposite directional configurations collapse onto identical values under cosine-based measures.

##### *3.2.* Decomposition of empirical phase synchronization (Fig. SF. 1b)

To show what each component carries in real data, we averaged the CVPS of every ICN pair across time and across the fBIRN cohort and visualized the population-level phase organization. The left matrix displays the mean phase of each pair, that is, the full directional structure. The right matrix displays the corresponding cosine (CRP) and sine (SRP) components in complementary triangles. The CRP component reflects only alignment strength, the same coupling magnitude that functional and dynamic functional connectivity quantify, whereas the SRP component carries the systematic lead-lag relationships between domains. The block structure visible in the SRP component and absent from the CRP component shows that meaningful organization resides in the discarded sign.

##### *3.3.* Single-connection illustrations in empirical data (Fig. SF. 1c, d)

To show that the directional component changes empirical conclusions at the level of individual connections, we examined two pairs of NeuroMark ICNs. For each subject, the mean CVPS was computed across the time series, and group differences between controls and individuals with schizophrenia were tested on the cosine of relative phase with a two-sample t-test and on the full directional phase with Watson's $U^2$ test. For the precuneus and thalamus pair (Fig. S1c), the cosine detected no group difference ($P = 0.78$) while the directional phase separated the groups ($P = 3.1 \times 10^{-5}$), a difference carried purely by timing. For the hippocampus and superior parietal lobule pair (Fig. S1d), the pattern reversed, with the cosine flagging a difference ($P = 6.4 \times 10^{-3}$) that the full phase distribution did not support ($P = 0.18$), indicating that magnitude-based comparisons can register differences in alignment strength that do not reflect a change in the underlying phase configuration. Retaining the full geometry therefore both reveals differences the cosine misses and guards against differences the cosine spuriously flags.

##### *3.4.* Phase estimation accuracy under non-stationary BOLD-like conditions (Fig. SF. 1e)

To compare Gabor wavelet against Hilbert phase estimation for non-stationary signals, we synthesized 1,000-sample (600 seconds) non-stationary traces under realistic BOLD conditions. A random frequency-modulated carrier was generated by integrating an instantaneous frequency that slowly wobbled between 0.035 Hz and 0.065 Hz (sinusoidal modulation at 0.01 Hz). The carrier was multiplied by a low-frequency amplitude envelope (0.004, 0.012, and 0.022 Hz components) and normalized, and this carrier was encoded into a wave signal with a non-stationary phase corresponding to the simulated frequency trajectory. Zero-mean Gaussian noise was added at signal-to-noise ratios of 50, 40, 30, 20, and 10 dB after band-passing (0.03 to 0.07 Hz, 0.018 to 0.098 Hz transition) to match BOLD spectral content. The instantaneous phase was extracted with the conventional Hilbert transform applied to the band-passed signal and with the complex Gabor wavelet applied directly to the raw trace $(f_0 = 0.05\,Hz, f_{bw} = 0.02\,Hz)$. Accuracy was quantified as the root mean square error (RMSE) against the known ground-truth phase trajectory, across 1000 independent noise realizations per SNR level, and the two estimators were compared with a paired test at each level.

#### Supplementary Fig. (SF) 1. Directional phase carries information that cosine-based measures discard, and the Gabor wavelet tracks it more accurately than the Hilbert transform.

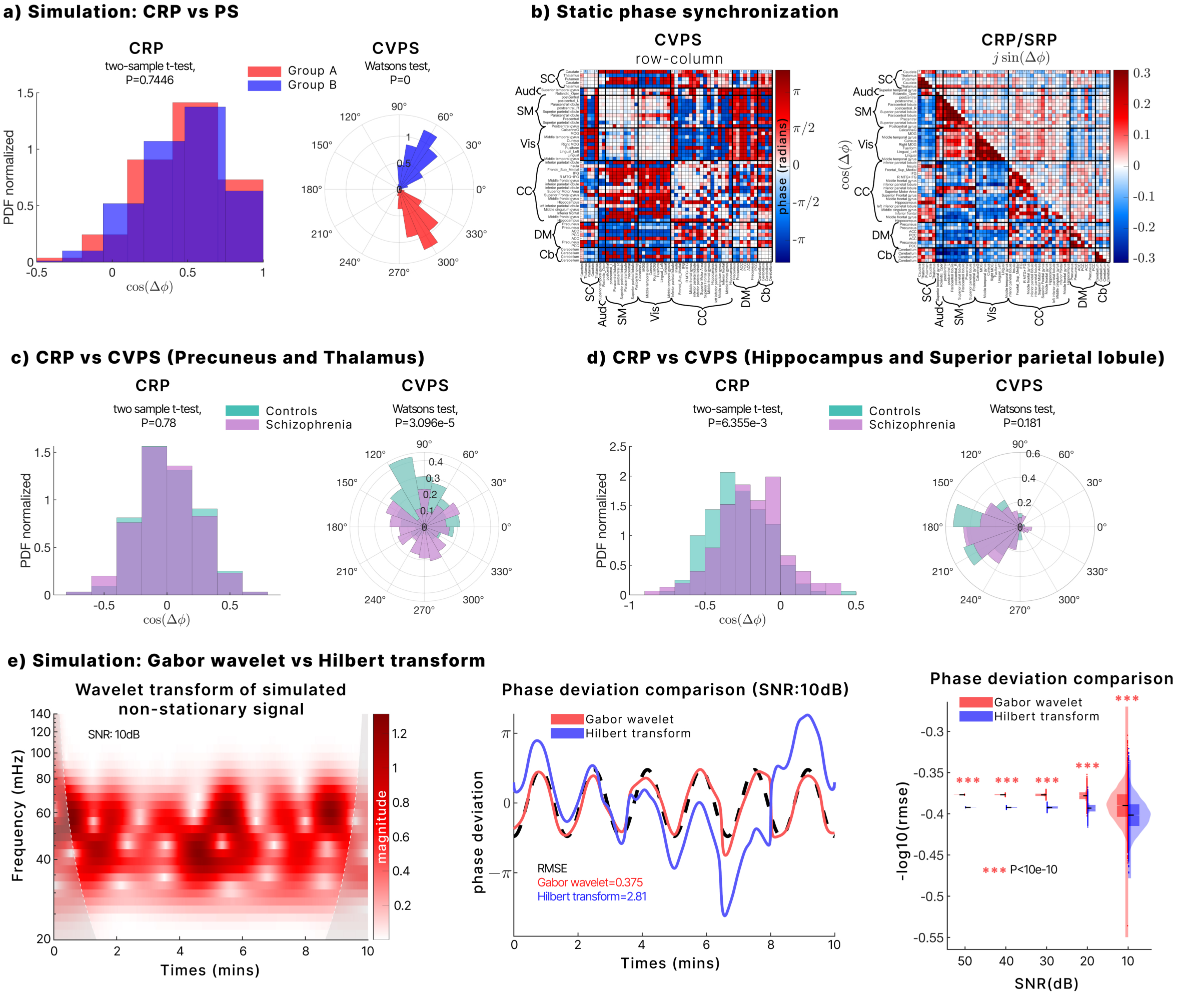


*(a) Two synthetic groups with mean phase offsets of +60° and –60° are indistinguishable using the cosine of the phase difference (two-sample t-test, P = 0.74) but clearly separable using the full circular phase (Watson's U², P ≈ 0), so opposite directional configurations collapse onto identical cosine values. (b) Population-level phase organization in fBIRN. Left, the average CVPS across the cohort, displaying the full phase of each ICN pair. Right, the corresponding cosine (CRP, $\cos\Delta\varphi$) and sine (SRP, $j\sin\Delta\varphi$) components, showing that the systematic lead-lag structure between domains resides in the sine component that magnitude-based measures discard. (c, d) Group comparisons at two example connections, testing the cosine with a two-sample t-test and the full phase with Watson's U². For precuneus and thalamus (c), the directional phase separates controls from patients (P = 3.1 × $10^{-5}$), whereas the cosine does not (P = 0.78). For the hippocampus and superior parietal lobule (d), the cosine flags a difference (P = 6.4 × $10^{-3}$) that the phase distribution does not support (P = 0.18). Direction both reveals and corrects. (e) Phase estimation under non-stationarity. Left, the wavelet transform of a simulated frequency-modulated signal (instantaneous frequency 0.035 to 0.065 Hz; SNR 10 dB). Middle, the Gabor estimate (red) tracks the ground-truth phase (black dashed), whereas the Hilbert estimate (blue) diverges (RMSE 0.375 versus 2.81). On the right, RMSE across SNR levels confirms that the Gabor wavelet is more accurate at every noise level (paired tests, *** $P < 10^{-4}$).*

### 4. Directional coordination regime estimation

#### *4.1.* K-means clustering

Whole-brain CVPS vectors were partitioned using k-means clustering to identify recurring coordination regimes. Each complex edge value $\Delta\varphi_{x \to y} = a + jb$ was “flattened” into the two real features $[a, b]$, giving a $2E$-dimensional real vector per TR ( $E =$ number of directed edges). Because $\Delta\varphi_{x \to y}$ is simply the sign-reversed complement of $\Delta\varphi_{y \to x}$, we retained one direction per pair to halve dimensionality; the omitted orientation can be recovered by sign inversion after clustering. This real-valued representation permits standard *k*-means without bespoke complex metrics while giving equal weight to cosine and sine components. Once the algorithm converged, each centroid was “unflattened” (recombined $a$ and $b$ as $a + jb$), producing a full complex-phase template for every recurring brain state. We implemented the *k*-means algorithm with k ranging from 1 to 10, set the maximum number of iterations to 10,000, and used 20 random initializations to ensure robust convergence.

The optimal number of clusters was selected as K = 3 using the elbow criterion (Supplementary Fig. 2a). To assess cluster-number sensitivity, we additionally examined K = 2 and K = 4 solutions. The K = 2 solution under-resolved the directional coordination structure by merging the feedback- and feedforward-like regimes, whereas the K = 4 solution preserved the feedback and feedforward regimes while subdividing the integrative regime into two related subtypes (Supplementary Fig. 2b–c). These analyses support K = 3 as the most parsimonious resolution for capturing the dominant feedback, feedforward, and integrative coordination architecture.

**Supplementary Fig. (SF) 2. Cluster-number sensitivity supports K = 3 as the parsimonious directional coordination solution.**

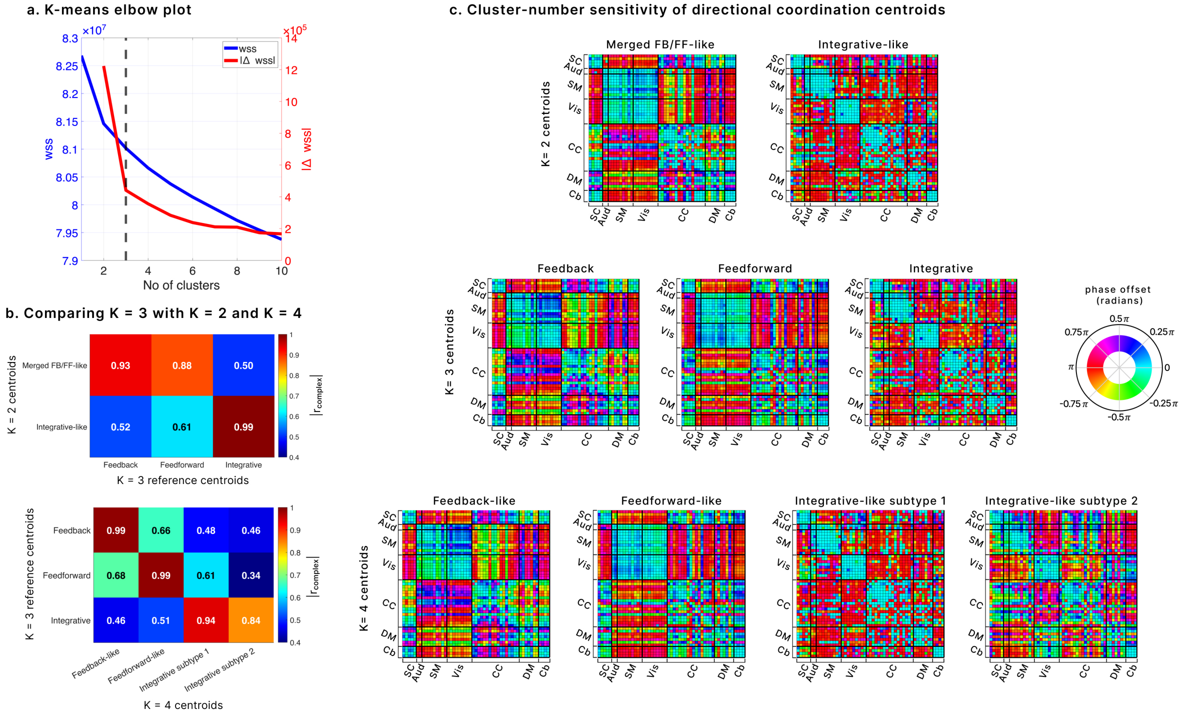


*(a) K-means elbow analysis shows a marked reduction in within-cluster sum of squares up to K = 3, supporting K = 3 as a parsimonious solution. (b) Absolute complex correlations between centroids across K values. The K = 2 solution under-resolves the K = 3 structure: one K = 2 centroid jointly matches the feedback and feedforward regimes, whereas the second closely matches the integrative regime. The K = 4 solution preserves the feedback and feedforward regimes while subdividing the integrative regime into two related subtypes. (c) Phase-offset centroid matrices for K = 2, K = 3, and K = 4 visually show the corresponding merge/split structure. Together, these analyses indicate that K = 3 captures the dominant feedback, feedforward, and integrative coordination architecture without collapsing distinct regimes or over-splitting them.*

#### *4.2.* K-means distance selection

When each phase is embedded on the unit circle as $v(\varphi) = [\cos\varphi, \sin\varphi]$, the squared Euclidean distance between two angles $\varphi_x$ and $\varphi_y$ is:

$$d_E^2(\varphi_x, \varphi_y) = \left(\sqrt{(\cos\varphi_x - \cos\varphi_y)^2 + (\sin\varphi_x - \sin\varphi_y)^2}\right)^2$$

$$d_E^2 = 2 - 2\cos(\varphi_x - \varphi_y) = 4\sin^2\left(\frac{\Delta\varphi}{2}\right), \qquad \Delta\varphi = |wrap_\pi(\varphi_x - \varphi_y)| \in [0, \pi] \qquad \text{(SE. 3)}$$

Because $4\sin^2\left(\frac{\Delta\varphi}{2}\right)$ increases strictly on $[0, \pi]$, minimizing the Euclidean distance is exactly equivalent to minimizing true angular separation $\Delta\varphi$; hence ordinary Euclidean k-means performs bona-fide angle clustering, and the arithmetic mean of the embedded points returns the circular mean.

For robustness in our very high-dimensional feature space (thousands of cosine/sine pairs), we also consider the Manhattan (city-block) distance

$$d_M(\varphi_x, \varphi_y) = |cos\,\Delta\varphi| + |sin\,\Delta\varphi| \qquad \textit{(SE. 4)}$$

Although $d_M$ is not an exact monotone transform of $\Delta\varphi$, it preserves the ranking of separations over most of $[0, \pi]$, and is less sensitive to distance concentration in high dimensions. Empirically, clustering solutions (state centroids and assignments) were nearly identical under both distances (SF 3); we report results with city-block due to improved numerical stability under high-dimensionality[15].

**Supplementary Fig. (SF) 3. Directional coordination centroids are robust across distance metrics.**

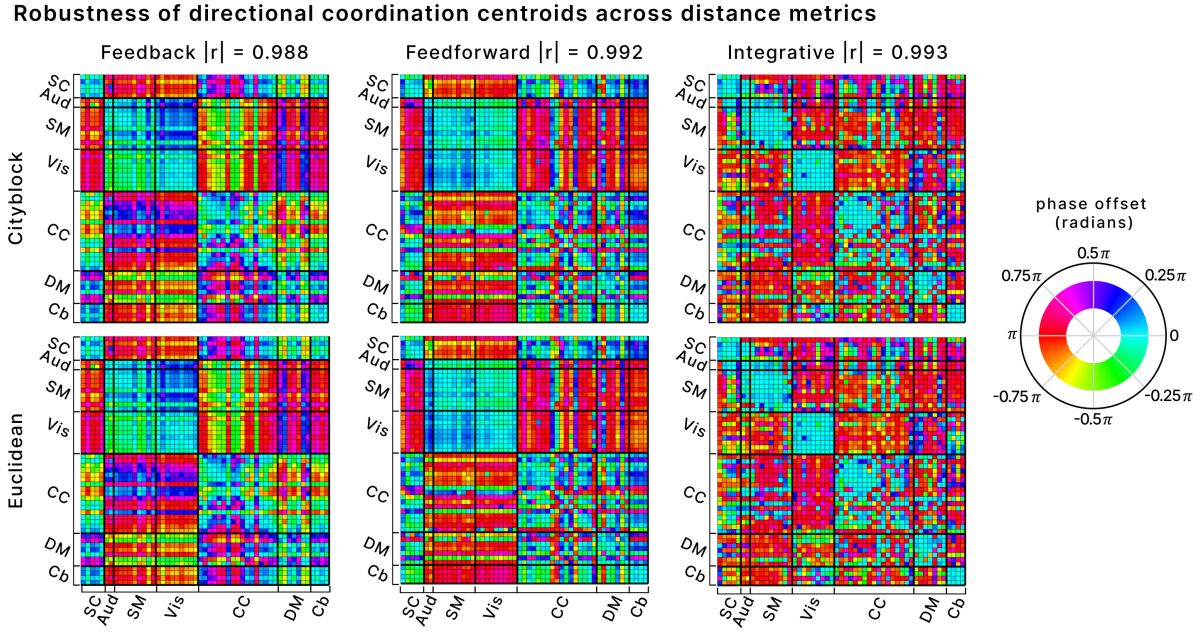


*Directional coordination regimes were estimated independently using cityblock and Euclidean distances in the embedded complex phase space. Centroids were reconstructed as complex-valued edge patterns and visualized by their phase offset in radians. Euclidean centroids were reordered to match cityblock centroids using one-to-one permutation matching that maximized absolute complex correlation. The matched centroid correlations were high for all three regimes: feedback,* $|r| = 0.988$*; feedforward,* $|r| = 0.992$*; and integrative,* $|r| = 0.993$*. These results indicate that the recovered directional coordination structure is stable across clustering distance metrics.*

### 5. Vascular and non-neural control details

For the regime-dependence control (Fig. 5b), the cosine and sine of relative phase were computed per pair and time point as $CRP = cos(\Delta\varphi)$ and $SRP = sin(\Delta\varphi)$, summarizing undirected coupling and signed lead-lag, respectively. At the group level, similarity between the feedback and feedforward regimes was quantified as the correlation across the 1,378 pairs of the CRP pattern and of the SRP pattern. At the single-subject level, we computed within each scan the percentage of pairs whose SRP sign differed between the two regimes.

For the frequency-dependence control (Fig. 5c), the group-mean signed relative phase was recomputed for every pair in six narrow sub-bands spanning the resting-state range, with center frequencies of 0.02, 0.035, 0.05, 0.07, 0.10, and 0.13 Hz, a half-bandwidth of 0.01 Hz, and filter edges trimmed. From these, we derived, per cohort and frequency, three quantities. Directional strength was computed as $|\langle sin\,\Delta\varphi\rangle|$, averaged over pairs and normalized to the 0.02 Hz band. The implied delay was computed as $\tau(f) = \frac{\Delta\varphi(f)}{2\pi f}$, where $\Delta\varphi(f)$ is the mean phase offset. A fixed conduction delay τ produces a phase offset $\Delta\varphi(f) = 2\pi f\tau$, predicting directional strength rising in proportion to f, a constant implied delay, and a frequency-flat directional pattern, whereas a genuine band-limited phase relationship predicts approximately flat strength, an implied delay falling as roughly 1/f, and a directional pattern that peaks in the analysis band. Directional pattern specificity was computed as the correlation across pairs between each sub-band's directional pattern and that of the 0.05 Hz analysis band. Curves were computed per subject, with figures showing cohort means and 95% confidence intervals across subjects.

At the lowest sub-band (0.02 Hz), the shorter scans contain approximately six cycles, so the low-frequency result is anchored on HCP Adult, whose longer acquisitions contain approximately seventeen cycles, where the same pattern holds. Cohort-specific preprocessing for these analyses is described in the dataset section above.

### 6. Directional driver scores

Node-level driver scores aggregate the edge-level information defined in the main text (Equation 3). For each edge connecting nodes i and j, the direction of the contribution was determined by the sign of the phase difference between the two nodes, so that the leading node receives a positive contribution and the lagging node a negative one. Each node's driver score was then computed by summing these signed contributions across all of its edges, weighted by the discriminative score of Equation 3, yielding each region's net directional drive within the regime. Positive driver scores therefore identify regions that tend to lead the rest of the brain in that regime, and negative scores identify regions that tend to lag; these are the values projected onto the cortical surfaces in Figs. 1c to 1e and 4c and entered into the directional hierarchy index and the gradient-alignment analyses. The delta-weighting applied before the gradient analyses, and its robustness, are described in the external hierarchy alignment section below.

### 7. External hierarchy alignment

The directional drivers align with the Margulies principal gradient under the exact Glasser-360 parcellation in both manipulations of the task and anesthesia cohort (Supplementary Fig. 4), with the robustness checks and convergent evidence detailed below.

#### *7.1.* Gradient projection into component space

The principal gradient was oriented using canonical sensory anchors (V1, A1, M1) against association anchors, so that higher values correspond to the association end of the hierarchy. Because the fBIRN regimes are defined on distributed NeuroMark components, the gradient was painted into MNI space via the Glasser atlas, resampled to the NeuroMark grid, and projected onto each component as the positive-weight mean over its cortical voxels, yielding gradient positions for 42 cortical components; subcortical and cerebellar components were excluded, as the gradient is defined for cortex only. For the task and anesthesia cohort, gradient values were assigned directly by Glasser parcel, spanning 329 cortical regions after exclusion of regions without network assignments.

##### *7.2.* Robustness to driver weighting and gradient kernel

The alignment between directional drivers and the Margulies principal gradient was robust to the two main analytical choices. Under the exact Glasser-360 parcellation (task and anesthesia cohort), the correlations were essentially unchanged whether or not driver scores were delta-weighted (feedforward rho = –0.83 weighted versus –0.825 unweighted; feedback +0.71 versus +0.710; integrative near zero in both), indicating that the primary result does not depend on this choice. In fBIRN, the feedback alignment held with or without weighting (rho = +0.76 weighted; +0.83 unweighted), whereas the feedforward alignment reached significance only with delta-weighting (rho = –0.64 versus –0.22 unweighted). This is expected: in the coarse 42-component NeuroMark space, the baseline lead-lag common to all regimes constitutes a larger fraction of each component's net driver, so isolating regime-specific directional structure requires down-weighting the shared component, whereas the finer 329-region Glasser parcellation renders this unnecessary. Repeating the analysis with a cosine-affinity kernel in place of the normalized-angle diffusion-map embedding gave essentially identical correlations, confirming the result is not specific to the gradient-estimation method.

##### *7.3.* Convergent cytoarchitectonic support

As a non-connectivity-based check in the task and anesthesia cohort, we compared directional drivers between cytoarchitectonically defined sensory and association classes (Mesulam laminar levels and von Economo types, from the atlas' own Glasser segmentation). The driver difference reversed by regime as predicted: feedforward association-minus-sensory = –8.3 (Mesulam) and –16.0 (von Economo); feedback = +9.6 and +15.2; integrative intermediate. This histology-based support holds at the sensory-versus-association dichotomy but not at the finer laminar ranking (paralimbic and limbic classes sit off the principal gradient), which is why the continuous connectivity gradient, rather than cytoarchitectonic class, is used as the primary external hierarchy in the main text.

##### *7.4.* Replication across both within-subject manipulations

The alignment reproduced under the exact Glasser-360 parcellation in both manipulations of the task and anesthesia cohort (Supplementary Fig. 4): motor and spatial imagery (feedforward rho = –0.79, feedback +0.76, integrative n.s.) and propofol-induced loss of consciousness (feedforward rho = –0.83, feedback +0.71, integrative n.s.). Both directional poles exceeded the spatial-autocorrelation-preserving null (Moran spectral randomization, 1,000 surrogates; p = 0.001) in each manipulation, and the integrative regime fell within the null in both. That the same alignment appears under two independent manipulations of the same subjects shows it is a property of the directional regimes rather than of any single condition.

**Supplementary Fig. (SF) 4. The directional axis aligns with an independent cortical hierarchy in both within-subject manipulations (Glasser-360).**

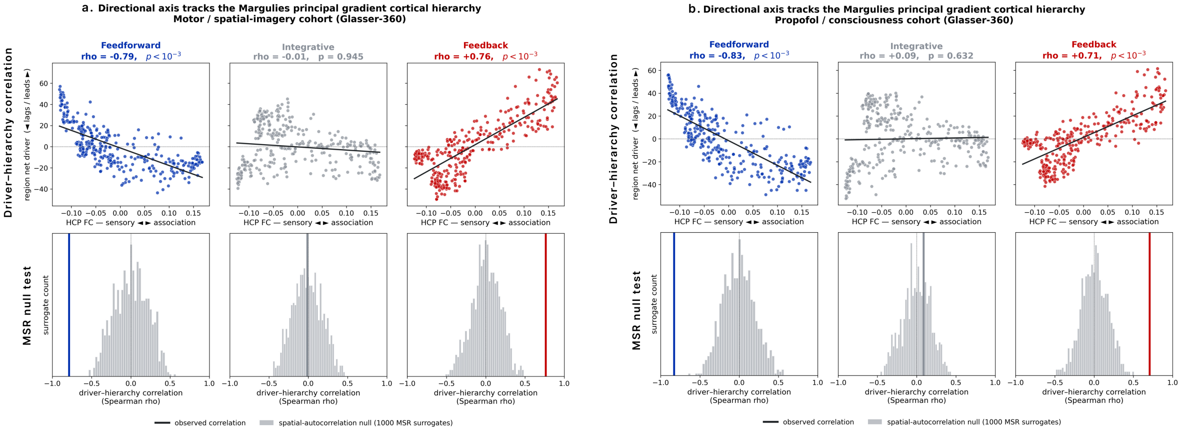


*Per-region net directional driver in each regime plotted against the Margulies principal functional-connectivity gradient, derived from an independent HCP cohort's connectivity and blind to our lead-lag and system grouping (sensory low, association high).* ***(a)*** *motor and spatial imagery;* ***(b)*** *propofol-induced loss of consciousness. In each, feedforward drivers correlate negatively with the gradient (sensory regions lead) and feedback positively (association regions lead), while the integrative regime shows no alignment. Bottom rows: observed correlation (colored line) against a spatial-autocorrelation-preserving null (1,000 Moran spectral-randomization surrogates, gray); both directional poles fall outside the null (p = 0.001) and the integrative regime within it. Effect sizes exceed those in fBIRN under the distributed NeuroMark parcellation (Fig. 2g), consistent with attenuation when a parcel-based gradient is projected onto spatially distributed components rather than assigned to discrete regions.*

### 8. Decoding feature sets

From the sampled time points, we extracted five feature sets, each operationalizing one organizing principle. (1) Directional coordination, the CVPS lead-lag features themselves, one signed relative phase per pair, approximately 1,378 features. Because the regimes are defined in this space, this set serves as a reference and upper bound on decodability and not as a competing principle. (2) Propagating-wave patterns, the real and imaginary coefficients of the leading complex-principal-component (traveling-wave) modes of the analytic signal (6 modes), capturing the instantaneous wave configuration. (3) Propagating-wave phase, the instantaneous phase of the dominant traveling-wave mode, entered as the cosine and sine of the leading mode phases, capturing the position within the wave cycle. (4) Global and arousal signal, the global mean signal, its temporal derivative, and its Hilbert phase entered as cosine and sine. (5) Principal gradient projection, each frame's projection onto the group principal functional-connectivity gradient, together with its temporal change. All features were z-scored within the training folds, and the identical sampling, folds, and classifier were applied to every feature set.

#### 9. Markov chain metrics, formal definitions

For each subject, the transition probability matrix P was estimated from the regime sequence, with entry $P(i,j)$ the number of observed transitions from regime $i$ to regime $j$ divided by the number of visits to regime $i$. Mean dwell time was computed empirically as the average length of uninterrupted runs in each regime, and fraction rate as the proportion of time points assigned to it. The stationary distribution $\pi$ was obtained as the leading left eigenvector of P, normalized to sum to one, and gives the long-run equilibrium occupancy implied by the transition structure. The spectral gap was computed as one minus the modulus of the second-largest eigenvalue of P, and the mixing time as the smallest number of steps t for which the total variation distance between the evolving distribution and $\pi$ falls below a tolerance of $10^{-3}$. The entropy rate was computed as $H = -\sum_i \pi_i \sum_j P(i,j) \log_2 P(i,j)$ and expressed as a percentage of log 3, the maximum for a three-regime system, which renders the choice of logarithm base immaterial. Full derivations and properties of these quantities are given in Wiafe et al.[16] and standard treatments[17].

#### 10. Persistence landscape construction

Each regime's stability was defined as the product of its stationary probability and its self-transition probability, and the three stability values were normalized to sum to one, preserving their relative structure. The three regimes were placed at the vertices of an equilateral triangle, feedback at (−1, 0), feedforward at (1, 0), and integrative at (0, √3), providing a minimal simplex representation of the three-regime system. A continuous surface was generated on a 300 × 300 grid spanning x ∈ [−1.4, 1.4] and y ∈ [−0.4, 2.2] by centering an isotropic Gaussian kernel (σ = 0.45) at each vertex with amplitude equal to the regime's normalized stability, summing the three kernels, and negating the sum so that deeper basins correspond to more stable regimes. Group landscapes were computed from each group's stationary distribution and transition matrix, and both surfaces were scaled by the absolute value of the global minimum across the two groups, so that basin depths are expressed in units of the deepest basin, and differences reflect relative regime stability and not arbitrary scaling. The group-difference surface (Fig. 2b) was computed as the control landscape minus the schizophrenia landscape.

**Supplementary Table ST. 1: Within-patient associations are unchanged when chlorpromazine-equivalent dose is included as a covariate**

| Outcome | Predictor | P_FDR (original) | P_FDR (+CPZ) |
|---|---|---|---|
| Attention vigilance | Feedback mean dwell time | 2.24e-2 | 2.29e-2 |
| Attention vigilance | Spectral gap | 1.38e-2 | 2.44e-2 |
| Composite cognition | Spectral gap | 3.56e-2 | 4.1e-2 |
| Composite cognition | Mixing time | 3.58e-2 | 4.15e-2 |
| Positive symptoms | INT→INT transition | 1.22e-2 | 1.65e-2 |
| Positive symptoms | FB→INT transition | 1.22e-2 | 1.65e-2 |
| Positive symptoms | INT→FF transition | 1.22e-2 | 1.65e-2 |

*Each association from Fig. 3b and 3c was re-estimated within patients with chlorpromazine-equivalent (CPZ) dose added to the generalized linear model alongside age, sex, scanning site, and mean framewise displacement. P_FDR denotes the FDR-adjusted p-value. Patients without a recorded dose were excluded from the adjusted models. FB, feedback; FF, feedforward; INT, integrative.*

### REFERENCES.

11. Larivière S, Paquola C, Park B-y, et al. The ENIGMA Toolbox: multiscale neural contextualization of multisite neuroimaging datasets. *Nature methods.* 2021;18(7):698-700.
12. Oppenheim AV. *Discrete-time signal processing.* Pearson Education India; 1999.
13. Li D, Li X, Cui D, Li Z. Phase synchronization with harmonic wavelet transform with application to neuronal populations. *Neurocomputing.* 2011;74(17):3389-3403.
14. Wiafe S-L, Soleimani N, Calhoun VD. Dynamic Co-Modulation (DyCoM): A Unified Operator Framework for Dynamic Connectivity in Neuroimaging. *bioRxiv.* 2026:2026.2001.2022.701132.
15. Aggarwal CC, Hinneburg A, Keim DA. On the surprising behavior of distance metrics in high dimensional space. Paper presented at: Database Theory—ICDT 2001: 8th International Conference London, UK, January 4–6, 2001 Proceedings 82001.
16. Wiafe S-L, Kinsey S, Soleimani N, et al. Aberrant recovery of timescale-aligned amplitude balance links to symptoms and cognition in schizophrenia. *Translational Psychiatry.* 2026;16(1):429.
17. Levin DA, Peres Y. *Markov chains and mixing times.* Vol 107: American Mathematical Soc.; 2017.
